# Bacteriophage pharmacodynamics studied in an in vitro pharmacokinetic model of infection

**DOI:** 10.1101/2025.08.11.669673

**Authors:** Marie Attwood, Pippa Griffin, Patryk Smorowinski, Alan Noel, Melissa Haines, Andrew Millard, Karen Adler, Martha Clokie, Alasdair Macgowan

## Abstract

**Background:** Bacteriophage therapy offers an alternative way to counter the menace of increasing antimicrobial resistance. Despite its use in clinical practice for many decades the basic tools to study the translational pharmacodynamics of phages are not available and it is recognised that lack of understanding of phage pharmacokinetic/dynamics (PK/PD) is a severe limitation in individual patient use and clinical trial design.

**Methods:** Traditional in vitro PK/PD evaluation tools were used to assess the antibacterial effect of single exposures of a bacteriophage cocktail against 4 strains of *E. coli* with potentially different patterns of response to phage. Initially, time-kill curves (TKC) were performed over 48hr and subsequently a dilutional in vitro model (IVM) was used to assess the antibacterial effects over 72hr.

**Results:** In TKC, the four *E. coli* strains showed different patterns of kill and regrowth when exposed to phage with two strains showing a sustained drop in bacterial viable count and two showing initial kill and regrowth. Using the IVM similar bacterial pharmacodynamic patterns were observed, and phage titre increased inversely but consistently with *E. coli* kill.

**Conclusions:** An In vitro dilutional model can be used to study the antibacterial effect of a phage cocktail on *E*.*coli* showing strain-to-strain variation in bacterial killing and bacteriophage titre. Such models can be used to provide more nuanced information on phage pharmacokinetics/dynamics and translationally useful information for dosing in humans.

## Introduction

Since the discovery of the first antibiotic classes there has been concerns over longevity of antibiotic efficacy mainly related to antibiotic misuse, and potential antibiotic resistance^1^. Many in vitro tests have been designed to assess the efficacy of antimicrobials such as disk diffusion, micro broth dilution, time-kill studies and synergy testing^2^. Whilst these tests aim to prevent incorrect antibiotic prescribing, they fail to assess bacterial-drug interactions from a dynamic drug concentration perspective. Time kill curves (TKC) can be used for semi-dynamic evaluations and are often performed when evaluating novel small molecules but are not routinely performed in a diagnostic microbiology laboratory. TKC can be used begin to understand the relationship of antibacterial effect to a pharmacodynamic index (PDI) however, are limited by nutrients, single dose exposure and employment of fixed antimicrobial concentrations^3^. In vitro modelling systems overcome these issues and have been employed to provide more extensive and robust evaluations, which have been progressively utilised for drug development ^4^. In vitro models, dilutional or hollow fibre both provide replenishment of nutrients, precise bacterial infectious dose, human pharmacokinetic simulations, more detailed bacterial population analysis and the ability to determine emergence of resistance. This data allows the accurate determination of the dominant PDI, and its magnitude which has good translatability with In vivo studies and informs optimum dosage and duration of antibiotic therapy ^5^.

Despite rigorous evaluations for novel antibiotics or combination antibiotic therapies, antibiotic resistance continues to be a serious concern globally ^6^. This can be attributed to many factors; overuse of antibiotics in an unregulated manner, antibiotics use based on price rather than suitability, poor or outdated prescribing patterns, increasing numbers of highly populated regions, substandard antibiotic manufacture, ease of travel, natural bacterial evolution, bacterial genetic mutation due to incorrect pharmacokinetic and pharmacodynamic (PKPD) exposures^7, 8, 9^. National actions plans (NAP) have been put in place to address this issue in many countries. The UK NAP (Confronting antimicrobial resistance 2024 to 2029) states that; we should avoid unintentional antibiotic use, if antibiotics are required, we should optimise exposure (using PK/PD principles, therapeutic drug monitoring, etc), we should innovate (potential new therapies) and we should continue to educate and practise antimicrobial stewardship practices.

Phage therapy are viruses which specially target bacteria and whilst this is not technically a new solution as it was largely replaced by antibiotic use after world war II ^10^, this type of therapy remains effective and has the potential be a significant strategy in the reduction of AMR. Personalised phage therapy has been employed with good success in locations such as Eliava Institute in Tbilisi, Georgia ^11^. Pre-defined phage therapy is a finished product which contains multiple phage stains which has been successfully used in agriculture and farming ^12^ but could be applied in medicine. More recently a large observational study of phage therapy has shown both efficacy and safety especially for the combination of bacteriophages and antimicrobials ^13^ and a randomised controlled trial in urinary tract infection showed non inferiority of phage therapy alone to standard care ^14^. Despite these clinical findings it is by no means clear which is the optimal way phages should be dosed in man. Here we describe the development of the PK/PD In vitro dilutional model for phage simulations in order to inform future optimum phage-antimicrobial dosing.

## Materials and Methods

### Bacteria and media

Bacterial strains: 4 clinical *E. coli* strains from the collection at Southmead Hospital were used, one fluoroquinolone sensitive strain (C1.15), one fluoroquinolone resistant strain (C1.24), one ESBL producing strain (C1.52) and one NDM producing strain (C1.68). The antibiograms of each strain are shown in Supplementary material Table S1. The medium used for PMIC determination (Attwood et al 2024) ^15^ was Mueller-Hinton broth II (BD 212322). Media used for phage propagation challenge strains was Mueller Hinton agar (Oxoid PO0152A).

### In vitro pharmacokinetic model

An in vitro one-compartment pharmacokinetic model (IVM, Electrolab, Tewkesbury, UK) was used to simulate the target phage concentration at site of infection (rather than phage oral dose titre), specifically mean free-phage concentrations associated with experimental doses of phage retrieved in poultry ^17^. The apparatus has been described many times before for antibiotic use (Noel et al 2024) ^18^. The configuration of the model was largely the same as described. The central culture chamber was inoculated to reach a bacterial concentration of 1.5 x10^6^ CFU/mL. This is connected to a main reservoir via silicone tubing to ensure nutrient addition/ mimic flow rates or phage administration. Flow rates are determined by the speed of a peristaltic pump. A second peristaltic pump was used which is connected to a waste vessel via silicone tubing to ensure the volume of the central chamber remains the same. The temperature was maintained at 37°C and the broth constantly agitated. Aliquots obtained from the central chamber at hourly timepoints 0 to 8 hr and then at every 24-hrs until the end of the simulation. In these simulations administration of phage was by bolus injection and all experiments were performed in triplicate.

### Phages and *E. coli* host strains

Phages: JK08, 113, UP17 and host *E. coli* strains MH10, B31 and EA2 were used. JK08, 113 and UP17 were combined in a cocktail at a ratio of 1:1:1

### Bacterial killing curves

Bacterial time kill curves (TKC) were performed in 10 mL universals providing sufficient nutrients to perform simulations of up to and including 48 hrs. Viable counts were determined using a spiral plater (Don Whitley, Shipley, UK). Aliquots were plated onto Mueller Hinton agar plates for determination of viable counts (CFU/mL). The minimum level of bacterial detection was 1.0 x10^2^ CFU/mL. The aliquots were taken hourly from timepoint 0 to 8 h and then at every 24-hour increment until the end of the simulation. All experiments were performed in triplicate (Noel et al 2022)^16^.

### Phage propagation and titration Phage propagation and titration

Challenge bacterial strains colonies were inoculated in Mueller Hinton agar (Oxoid PO0152A) and grown overnight at 37°C aerobically. To prepare liquid culture, challenge strain colonies were inoculated into LB broth (Oxoid L3147) and grown overnight at 37°C agitated at 100 rpm. Phage propagation was performed by combining phages JK08, 113 and UP17 to the appropriate challenge strain at 1.0×10^7^ PFU/mL. Bacteria/phage mix cultures were incubated at 37°C at 100 rpm for 6.0 ± 1 hours. Bacteria/phage mix are centrifuged for 15 mins at 4200xg, and supernatant filtered using (0.2 micron pore size filters). Phage titre was determined by serial dilution (using SM buffer) and plated via plaque assay techniques. Phage titre stored at 4-8°C until use and appropriate dilutions performed as required.

### Phage direct spot testing

Bacterial cultures of appropriate challenge strains were grown overnight and then diluted 1/100 in LB and grown for 2 hrs to 0D550 of 0.2. 500 µL of the culture was added to 8 mL 0.5% (w/v) LB agar kept molten at 55°C and poured onto MHA plates. Plates (double agar overlay assay plates)were left to set prior to testing aliquots from the central chamber. TKC or IVM aliquots are serially diluted to determine PFU/mL (testing series). Once serial dilution has been performed 10 µL of each testing series is spotted directly onto previously made double agar overlay plaque assays. The plates were incubated overnight at 37°C and phage plaques enumerated.

### Data analysis

The antibacterial effect of each strain was described by the area under the bacterial kill curve (AUBKC) which was calculated according to the trapezoidal rule, to compare antibacterial effects. The data was plotted using the software package Graph Pad Prism Version 9 (San Diego,CA, USA).

## Results

### Phage minimum inhibitory concentrations

The phage cocktail produced the following PMIC’s: *E. coli* C1.15 = 1.0 x10^4^ PFU/mL, C1.24 = 1.0 × 10^6^ PFU/mL, C1.52 = 1.0 x10^5^ PFU/mL and C1.68 = 1.0 × 10^5^. The ranking of activity against the strains was C1.15 (cocktail most active), C1.52 and C1.68, then C1.24 (cocktail least active)

### Time Kill curve

Figure 1 shows time-kill curves with a phage - bacteria ratio of 1:1. Whilst the cocktail showed activity against all four strains, the pattern of bacterial kill over time was different. All strains showed initial killing with a 1–3 log drop in viable bacterial counts over the first 3 hrs, however only two strains (C1.15 and C1.52) displayed further reduction in CFU/mL with a 4-log drop at 6 hr and suppression of bacterial counts below the initial inoculum until 48 hrs. Strain C1.24 and C1.68 showed regrowth from 3 hours and bacterial counts were greater than the initial inoculum at 24 and 48hrs. Areas-under-the-bacterial-kill-curves (AUBKC) were calculated and are shown in Table 2. The ranking of cocktail activity against the strains was C1.15 (cocktail most active), C1.52, C1.24, then C1.68 (cocktail least active)

**Table 1.**
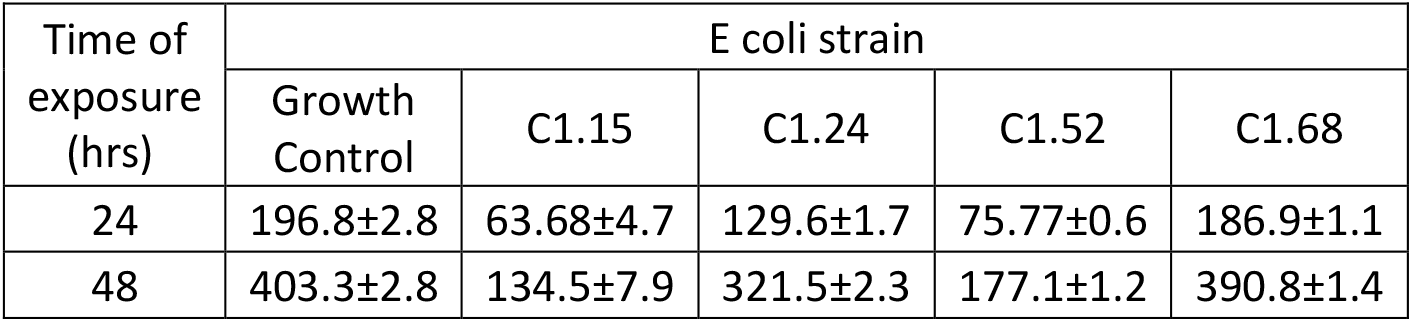
Antibacterial activity of the phage cocktail as measured by the area-under-the-bacterial-kill curve against *E coli*

| Time of exposure (hrs) | E coli strain |  |  |  |  |
| --- | --- | --- | --- | --- | --- |
|  | Growth Control | C1.15 | C1.24 | C1.52 | C1.68 |
| 24 | 196.8±2.8 | 63.68±4.7 | 129.6±1.7 | 75.77±0.6 | 186.9±1.1 |
| 48 | 403.3±2.8 | 134.5±7.9 | 321.5±2.3 | 177.1±1.2 | 390.8±1.4 |

**Figure 1.**
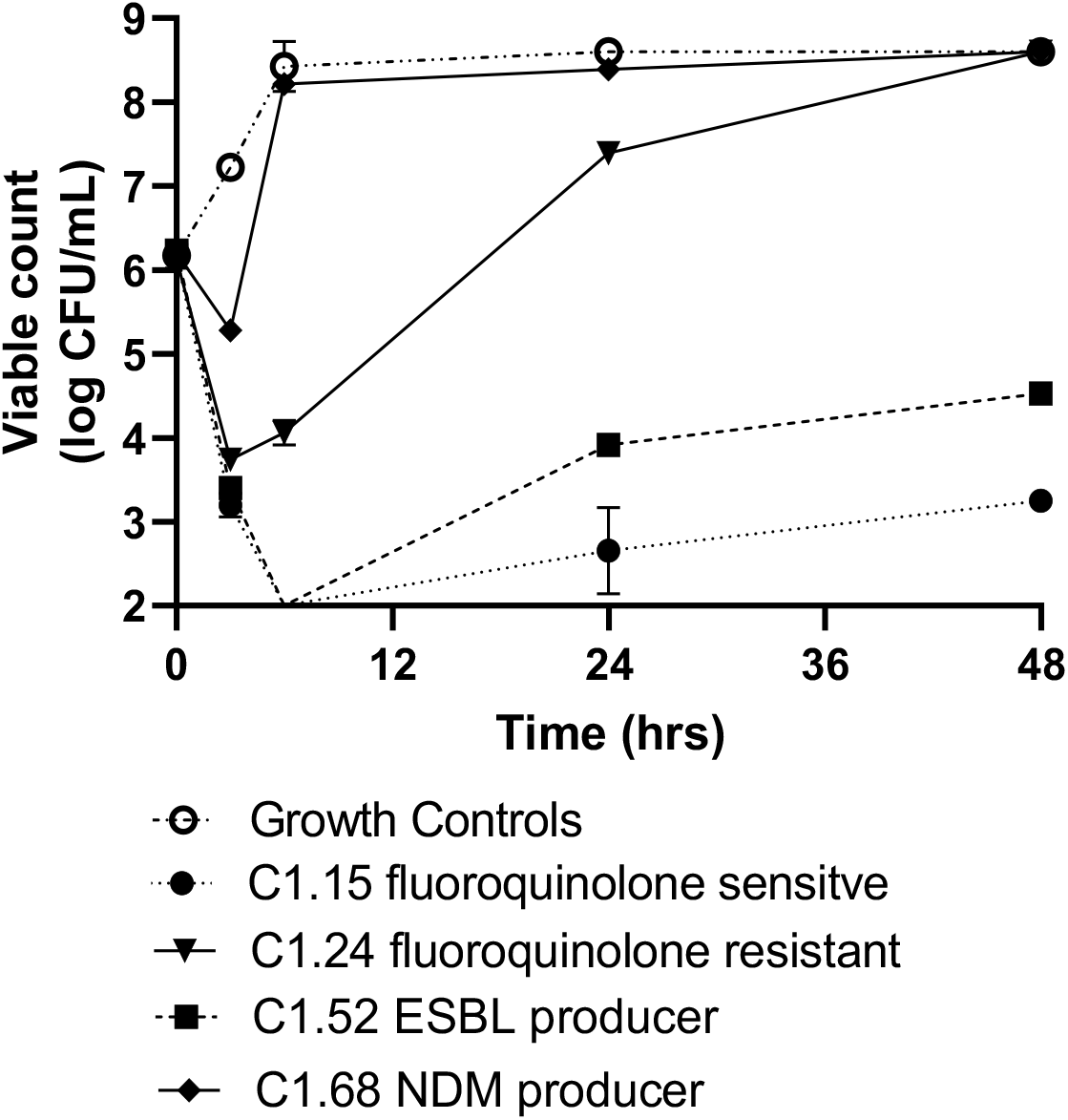
Antibacterial activity of the JK08,113, UP17 phage cocktail against four *E coli* stains

### In vitro model

Figure 2 shows growth curves of the four *E. coli* strains exposed to a phage cocktail dose of 1.5×10^2^ PFU/mL, bacterial inoculation 1.5×10^6^ CFU/mL. Unsurprisingly the IVM model data mirrors the TKC bacterial pharmacodynamics with one phage dose. C1.15 and C1.52 displayed >4 log reduction within the first 6 h of exposure, followed by bacterial regrowth to a bacterial load of approximately the initial inoculum by 72 h (Figure 2). With *E coli* C1.15 and C1.52 phage titres increasing significantly I in the first 1-2 hrs, in which the inundation point (optimum phage titre to bacterial CFU ratio) was crossed (phage titre PFU exceeded bacterial CFU/mL) after1-2 hr exposure. C1.24 showed a gradual reduction in CFU with a 1-2 log drop by 4 hrs, bacterial burden then increased until the end of the simulation. Inundation point (C1.24) was reached at 3 hrs with phage titre increasing gradually until 4 hrs. C1.68 showed minimal bacterial reduction with phage titre remaining low throughout the simulation.

**Figure 2.**
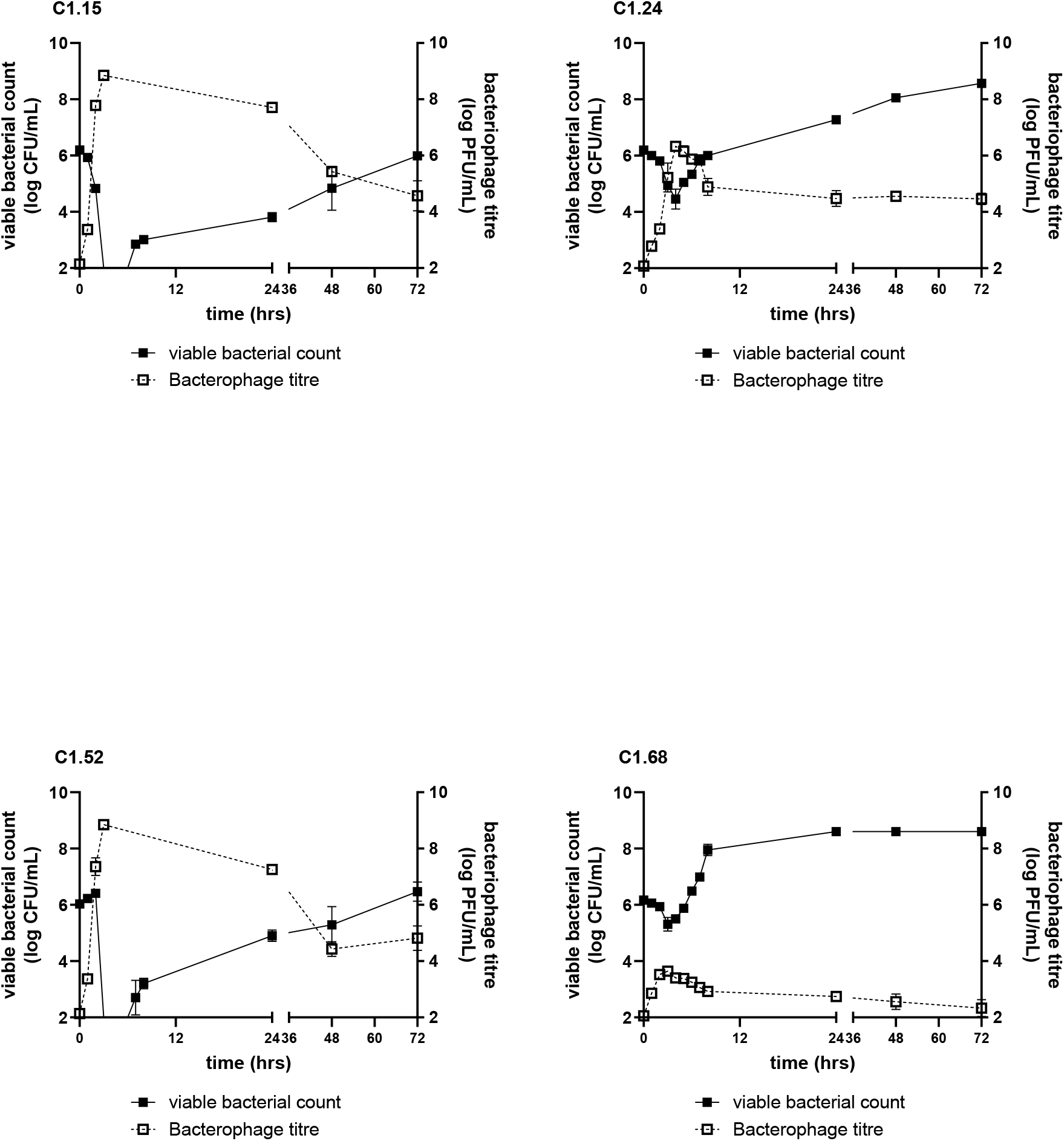
Antibacterial effect of a single exposure of bacteriophage cocktail on *E. coli* strains C1.15, C1.24, C1.52 and C1.68

Figure 3 shows the relationship between the area-under-the-bacteriophage concentration-time curve as a measure of total bacteriophage exposure and the AUBKC as a measure of bacterial kill. At all three time points analysed (24 h, 48 hrs or 72 hrs), as the phage exposure increased the bacterial kill increased. The ranking of cocktail activity against the strains in the IVM was C1.15 (cocktail most active, AUBKC at 24hrs 32.8±1.4), C1.52 (AUBKC at 24h 44.8±3.4), C1.24 (AUBKC at 24hrs 101.9±1.2), then C1.68 (AUBKC at 24hrs 133.6±2.0, cocktail least active).

**Figure 3.**
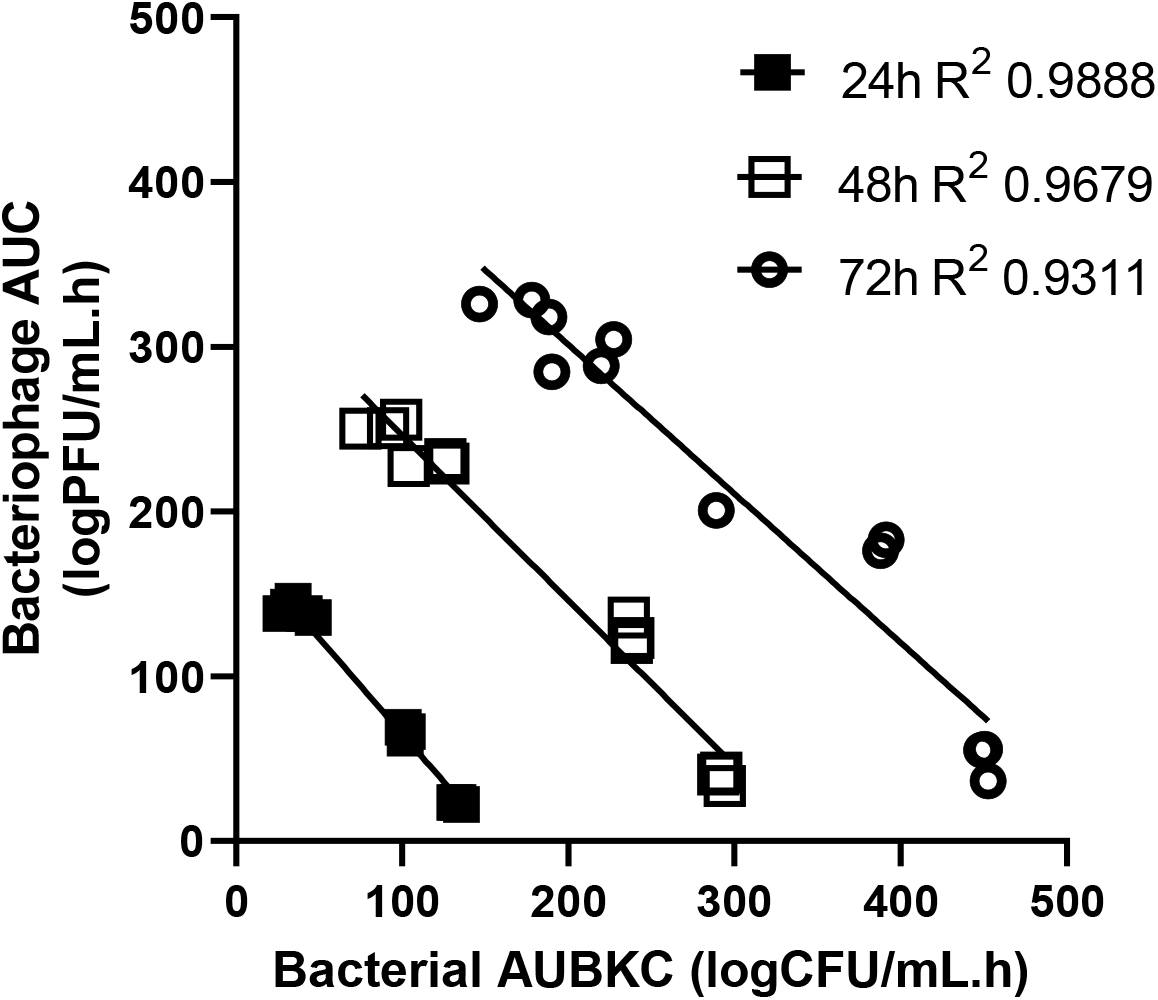
Relationship between antibacterial effect measured by AUBKC and bacteriophage exposure measured by the area under the bacteriophage titre time curve.

## Discussion

The in vitro pharmacokinetic/dynamic methodologies developed for small molecule antimicrobial evaluations have been shown to be an effective tool to assess phage antibacterial effects in this study and support previous work (Attwood et al, 2024)^15^. The results from time kill curves performed with a single phage exposures were in close agreement with similar single dose exposures in our in vitro pharmacokinetic model. The PMIC methodology may be useful for determining basic phage efficacy allowing the nuances of the phage-pathogen interaction to be initially assessed prior to performance of potentially more translatable experiments using time-kill curves and finally in vitro models. These approaches are essential in the design of phage dosing regimens for clinical trials as well as understanding phage-antibiotic interactions. A recent narrative review by Nang et al, 2023, highlighted the key role of further phage PK/PD in future phage patient therapy ^19^.

There are concerns that phage could not be studied within existing in vitro systems, specifically hollow fibre systems(HFIM). HFIM is a system of pumps, tubing and microfibers which allows for in vitro assessment of anti-infective compounds which is mimicking human conditions. These systems are used widely in Europe for PKPD evaluations however there are concerns remain for their use in phage evaluations due to the microfiber component. Fibers within the HFIM could; a) damage the phage tail fibres preventing the phage form being able to infect their hosts or b) phage may become bound to the membranes with the central compartment of the system. Whilst we acknowledge that this can be overcome by selecting appropriate cartridges (where materials, pore size and permeability), this is often specific and incur significant additional expense to each phage or phage cocktail evaluation. Dilutional in vitro models offer a solution to possible phage damage, binding (in that there are no fibres present) and a reduced expense (dilutional models are not single use). Washout of bacteria and phages is a potential source of concern with dilutional models but has been shown to be vastly overstated as a technical issue in antibacterial small molecule pharmacokinetic/dynamic studies ^20^.

We can state that this phage cocktail does not appear to bind to the dilutional in vitro modelling system components in either bolus or infusion administration. It appears that washout does not appear to be a factor either as various target phage titres used as targets 10^6^, 10^4^ and 10^2^ (PFU/mL) with excellent recovery of phage within standard IVM tolerance limits, data displayed in Figure 4 and 5.

**Figure 4.**
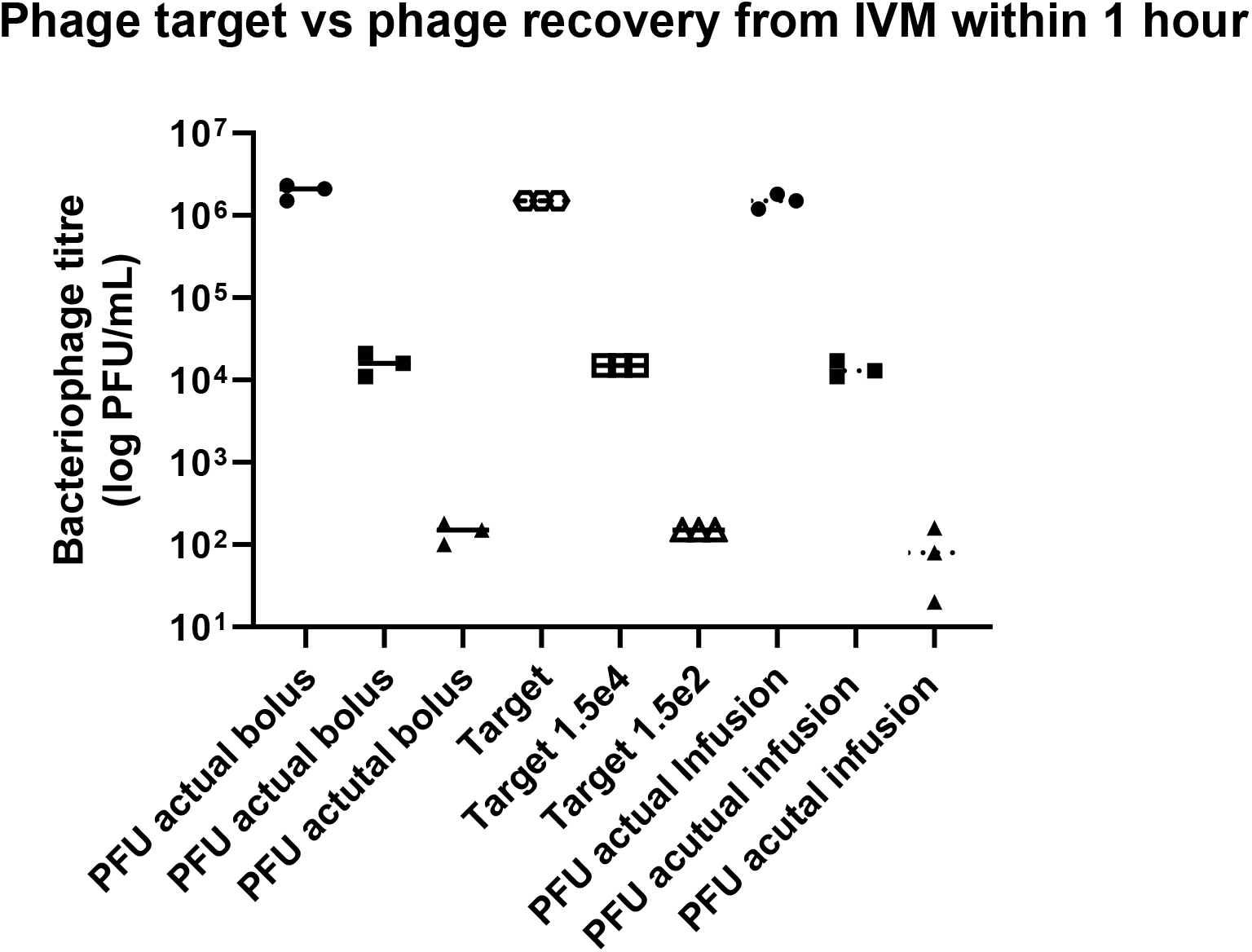
Assessment of phage binding with materials within the IVM system. Phage target dose (using corresponding bacterial host) vs actual phage titre recovery within 1 hour.

**Figure 5.**
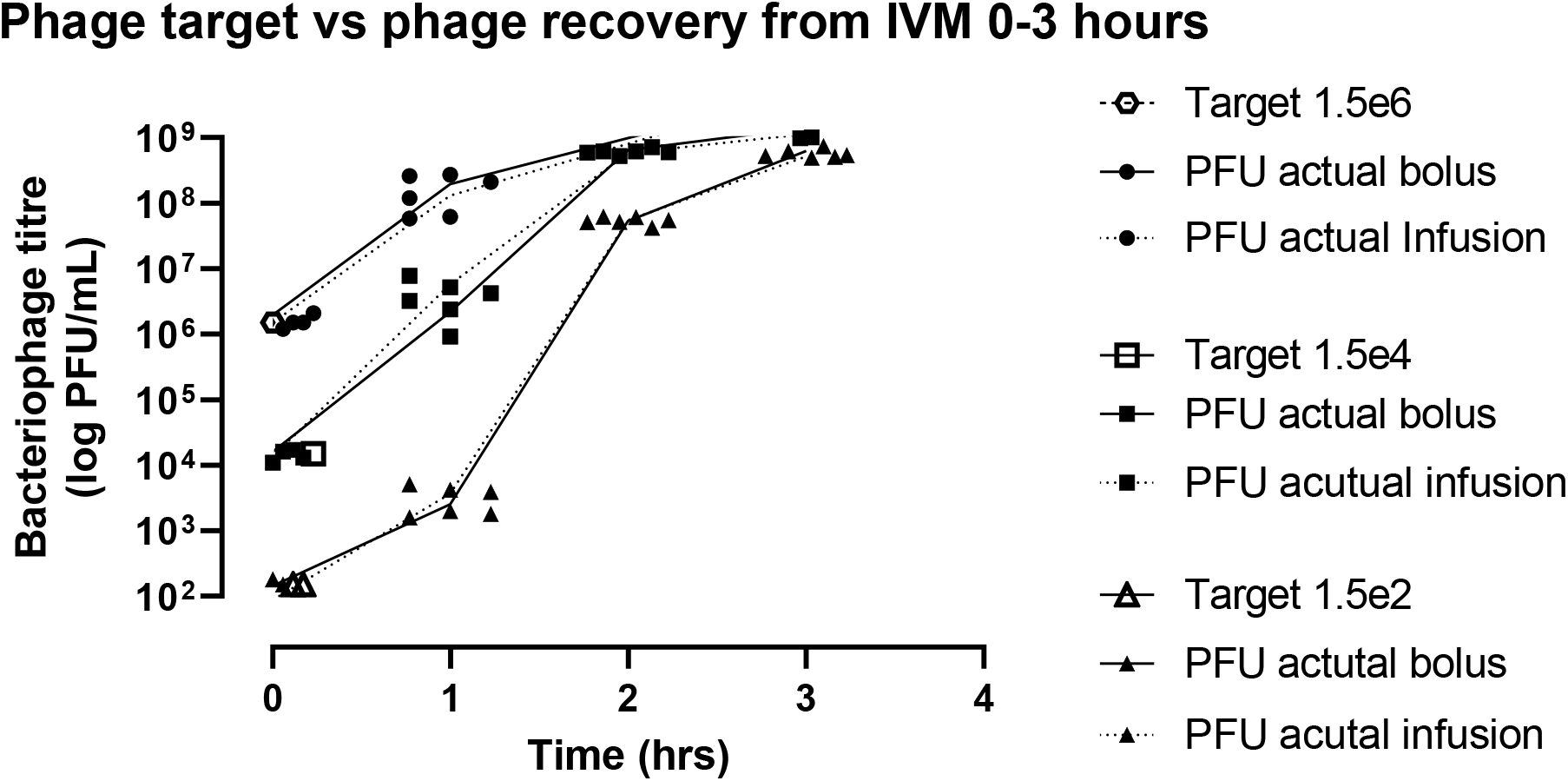
Assessment of phage binding with materials within the IVM system. Phage dose (using corresponding bacterial host) vs phage titre recovery within 3 hours.

Further work should now be performed, such as, phage dose/exposure escalation studies as well as on the potential timing and effect of second and subsequent phage exposures. This will enable the effect of phage dosing regimens on bacterial kill, regrowth and resistance to phage to be explored and more translationally useful information to be acquired.

Dilutional in vitro models also allow performance of extended simulations over up to 14 days which will help determine optimum phage - bacterial ratio, emergence of resistance of phage (from a phenotypic and genetic perspective), effect of different administration routes, effectiveness of multiple phage doses and combination of phage with antibiotics to be studied in conditions more akin to human therapy.

Whilst there is pharmacokinetic information available for specific phage cocktails, there is a paucity of phage pharmacokinetic data at target site concentrations ^21^. If this data becomes available, it will then be possible to mimic these using in vitro models to accurately simulate appropriate half-lives or addition of additional phages as required to mimic phage proliferation. Although it could be argued that perhaps pharmacokinetics at target site is not required, if all successful phage cocktails proliferate at a predictable and consistent rate, which are proportional and dependant on burst size. To determine this, we can use of these IVMs to answer such questions.

Whilst acknowledging that there is a great deal of development work left to do, it seems that dilutional in vitro models will be an essential element in determining optimum phage dose, dosing frequency and duration for use in humans.

## Transparency Declarations

MA and APM holds research grants/activities with Merck, Shionogi, InfectoPharm, GSK, Roche, BioVersys, VenatoRx Pharmaceuticals, iFAST, Technomede, Oxford Drug Design, JPIAMR and NIHR; APM provides consultancy advice to Shionogi, Roche, Bicycle Therapeutics and BioVersys.

## References

1. Muteeb, G., Rehman, M. T., Shahwan, M. & Aatif, M. Origin of Antibiotics and Antibiotic Resistance, and Their Impacts on Drug Development: A Narrative Review. Pharmaceuticals 16, 1615 (2023).

2. Balouiri, M., Sadiki, M. & Ibnsouda, S. K. Methods for in vitro evaluating antimicrobial activity: A review. J Pharm Anal 6, 71–79 (2016).

3. Foerster, S., Unemo, M., Hathaway, L. J., Low, N. & Althaus, C. L. Time-kill curve analysis and pharmacodynamic modelling for in vitro evaluation of antimicrobials against Neisseria gonorrhoeae. BMC Microbiol 16, 216 (2016).

4. Velkov, T., Roberts, K. D., Nation, R. L., Thompson, P. E. & Li, J. Pharmacology of Polymyxins: New Insights into an ‘Old’ Class of Antibiotics. Future Microbiol 8, 711–724 (2013).

5. Bulitta, J. B. et al. Generating Robust and Informative Nonclinical In Vitro and In Vivo Bacterial Infection Model Efficacy Data To Support Translation to Humans. Antimicrob Agents Chemother 63, (2019).

6. Ahmed, S. K. et al. Antimicrobial resistance: Impacts, challenges, and future prospects. Journal of Medicine, Surgery, and Public Health 2, 100081 (2024).

7. Otaigbe, I. I. & Elikwu, C. J. Drivers of inappropriate antibiotic use in low- and middle-income countries. JAC Antimicrob Resist 5, (2023).

8. Ehsan, H. Antibiotic Resistance in Developing Countries: Emerging Threats and Policy Responses. Public Health Challenges 4, (2025).

9. Zay Ya, K., Patel, J. & Fink, G. Assessing the impact of antimicrobial resistance policies on antibiotic use and antimicrobial resistance-associated mortality in children and adults in low and middle-income countries: a global analysis. BMJ Public Health 3, e000511 (2025).

10. Sahoo, K. & Meshram, S. The Evolution of Phage Therapy: A Comprehensive Review of Current Applications and Future Innovations. Cureus (2024) doi:10.7759/cureus.70414.

11. Parfitt, S. A. et al. The earliest record of human activity in northern Europe. Nature 438, 1008–1012 (2005).

12. Svircev, A., Roach, D. & Castle, A. Framing the Future with Bacteriophages in Agriculture. Viruses 10, (2018).

13. Pirnay, J.-P. et al. Personalized bacteriophage therapy outcomes for 100 consecutive cases: a multicentre, multinational, retrospective observational study. Nat Microbiol 9, 1434–1453 (2024).

14. Leitner, L., Kessler, T. M. & McCallin, S. E. Innovations in Phage Therapy for Urinary Tract Infection. Eur Urol Focus 10, 722–725 (2024).

15. Attwood, M. et al. Development of antibacterial drug + bacteriophage combination assays. JAC Antimicrob Resist 6, (2024).

16. Noel, A. R., Attwood, M., Bowker, K. E., MacGowan, A. P. & Albur, M. Comparative bactericidal activity of representative β-lactams against Enterobacterales, Acinetobacter baumannii and Pseudomonas aeruginosa. Journal of Antimicrobial Chemotherapy 77, 1306–1312 (2022).

17. Lorenzo-Rebenaque, L., Malik, D. J., Catalá-Gregori, P., Marin, C. & Sevilla-Navarro, S. In Vitro and In Vivo Gastrointestinal Survival of Non-Encapsulated and Microencapsulated Salmonella Bacteriophages: Implications for Bacteriophage Therapy in Poultry. Pharmaceuticals 14, 434 (2021).

18. Noel, A. R., Attwood, M., Bowker, K. E. & MacGowan, A. P. Pharmacodynamics of taniborbactam in combination with cefepime studied in an in vitro model of infection. Int J Antimicrob Agents 64, 107304 (2024).

19. Nang, S. C. et al. Pharmacokinetics/pharmacodynamics of phage therapy: a major hurdle to clinical translation. Clinical Microbiology and Infection 29, 702–709 (2023).

20. Dalhoff, A., Bowker, K. & MacGowan, A. Comparative evaluation of eight in vitro pharmacodynamic models of infection: Activity of moxifloxacin against Escherichia coli and Streptococcus pneumoniae as an exemplary example. Int J Antimicrob Agents 55, 105809 (2020).

21. Kang, D., Bagchi, D. & Chen, I. A. Pharmacokinetics and Biodistribution of Phages and their Current Applications in Antimicrobial Therapy. Adv Ther (Weinh) 7, (2024).

